# Superhydrophobic Sand Mulch and Date Palm Biochar Dramatically Boost Growth of *Moringa oleifera* in Sandy Soil: Insights into Evapotranspiration Budgeting and Metabolomic Profiling

**DOI:** 10.1101/2024.05.04.592506

**Authors:** Kennedy Odokonyero, Bob Vernooij, Batool Albar, Lisa Oki Exposito, Aishah Alsamdani, Amin Akhter Ghulam Haider, Nayara Vivian Huve Musskopf, Adair Gallo, Najeh Kharbatia, Himanshu Mishra

## Abstract

In response to the challenge of nutrient-deficient sandy soils and water scarcity due to excessive evaporative water loss in arid regions, we developed and tested two complementary soil amendment technologies: Superhydrophobic sand (SHS) mulch and an enriched date palm biochar. In a greenhouse pot experiment, we investigated the stand-alone and synergistic effects of SHS mulch (10 mm-thickness) and biochar (2% w/w) on *Moringa oleifera* plants under normal (**N,** 100% field capacity) and reduced (**R**, 50% of **N**) irrigation scenarios. Under **N** and **R**, SHS mulch reduced evaporation by 71% and 64%, respectively; while SHS+biochar reduced evaporation by 61% and 47%, respectively, in comparison with the control (*p* < 0.05). Total transpiration significantly increased in SHS plants by 311% and 385% under **N** and **R**, respectively. Compared with the control, transpiration increased in biochar-amended plants by 103% and 110%; whereas, its combination with SHS increased transpiration by 288% and 301% under **N** and **R**, respectively (*p* < 0.05). Irrespective of the irrigation regimes, we found superior effects of SHS, biochar, and their combination on plant height (62–140%), trunk diameter (52–91%), leaf area index (57–145%), leaf chlorophyll content index (11–19%), stomatal conductance (51–175%), as well as shoot (390–1271%) and root (52–142%) biomass, in comparison with the controls. Metabolomics analysis showed significantly higher relative abundance of amino acids, sugars, fatty acids, and organic acids in the leaves of control plants relative to other treatments, as a response to water or nutrient stress induced by excessive water loss through evaporation. Next, we found a higher concentration of D-Mannose, D-Fructose, glucose, and malic acid in plants grown with SHS or biochar treatment under **N** and **R** irrigation, attributed to increased water/nutrient-use efficiency and carbon assimilation because of higher photosynthesis rates than in the control plants. Our results show that, our complementary technologies could address the challenge of water loss via evaporation from soil and maximize soil nutrient retention for improving plant growth in arid regions. This could underscore the success and sustainability of irrigated agriculture and greening efforts in arid lands.

## INTRODUCTION

The use of plants as medicine for human health has been known for ∼60,000 years ago since the mid-Paleolithic age^1^, and yet only about 6% of the 391,000 vascular plants identified to date have been screened for their biological activity^2^ and 15% for their phytoconstituents^3^. Among those, *Moringa* constitutes an important genus with outstanding economic importance; used in traditional medicine and pharmacological screening.

The health-improving properties of *Moringa* is attributed to the numerous compounds produced by the plant as primary and secondary metabolites. Primary metabolites (e.g., carbohydrates, proteins, vitamins, nucleic acid, lipids, and hormones) are essential for plant development, growth, and reproduction while secondary metabolites (e.g. flavonoids, phenolic acids, isothiocyanates, tannins and saponins) are derivatives of the primary metabolites^4–6^. A growing spectrum of such therapeutic compounds in the plant makes it suitable in the treatment of oxidative stress, liver disease, neurological disease, hyperglycemia and cancer^7^. Regulated by the environmental conditions, the synthesis of such metabolites enable the plants to survive and protect themselves from abiotic and biotic stresses.

Of all the 13 species of *Moringa*, studies have mainly concentrated on *Moringa oleifera,* commonly grown in many African and Asian countries, for their food and medicinal uses^8^. However, of recent, *M. peregrina,* is another species that is becoming of importance for its traditional, nutritional, industrial and medicinal values in the Red Sea/Arabian peninsula and Northeast Africa ^9, 10^; and mostly in the South and North- Western regions of Saudi Arabia^11^. For instance, Saudi Al-Ula Governorate recently signed an ambitious agreement with *Dior Company* aimed at planting millions of *M. peregrina* trees in the region for the production of perfumes and cosmetics^12^.

Despite the economic potential that the cultivation of *Moringa* presents to the region, the challenges of water scarcity, owing to low precipitation, high temperature/heat stress, excessive water evaporation, and poor desert soils^13, 14^ hampers initiatives for large-scale cultivation of *Moringa* to tap in its pharmaceutical/medicinal benefits, restore vegetation cover and improve carbon capture, and biodiversity in the arid regions of Saudi Arabia. In response, plastic mulches could be deployed to enhance irrigation efficiency, as commonly practiced for crop production in arid regions; however, their non-biodegradability and disposal challenges render them unsustainable^15, 16^. Also, despite the excess fertilizers applied, the poor arid soil conditions with high infiltration exacerbate nutrient losses via percolation.

Due to biochar’s adsorptive properties and high porosity^17^, it has been used alongside other mulching technologies, as soil amendments for improving soil fertility, retaining nutrients and water in the soil, improving plant growth and productivity, as well as carbon sequestration in both managed and natural ecosystems^18–21^. Although afforestation efforts are increasing for the restoration of degraded arid lands under limitations of poor soil and water scarcity^22^, there is little information regarding the use of biochar and mulching for tree plantation^21^.

In an attempt to address the dual challenges of water scarcity and soil fertility for improving the cultivation of *Moringa* plants in Saudi Arabia, we have developed two complementary technologies, Superhydrophobic sand (SHS) mulch and biochar. SHS is a water-repellent material comprising of sand grains coated with thin layer of paraffin wax. Applying a 0.5–1 cm-thick SandX layer as a mulch can reduce evaporative water loss from the soil by 50-80%; plants transpire more and withstand the water/heat stress, and grow better in arid environments^23, 24^. The biochar used is a nutrient-enriched biochar derived from the pyrolysis of date palm biomass as feedstock. We have systematically optimized our biochar formulations for enhanced cation and anion exchange capacity, making them able to act as slow-release fertilizers for the essential macro and micronutrients; thus, can hold nutrients and prevent nutrient leaching in the soil.

Therefore, under the constraints of water scarcity from excessive evaporation and nutrient-deficient sandy soils, we evaluated the stand-alone and synergistic effects of our complementary soil amendment technologies on the growth of *M. oleifera* under normal and reduced irrigation conditions. We hypothesized that these bioinspired materials boost irrigation water and nutrient-use efficiencies towards improved plant growth in arid lands.

## Materials and Methods

### SHS mulch

The SHS used was produced using our lab-scale batch reactor by direct melting of paraffin wax via a heating process, which obviated the use of organic solvent as reported in our previous protocol^23–25^. A known mass of paraffin wax mixed with a known mass of sand (1:600 ratio) was heated in a rotating cement mixer for about 30 minutes until the surface of the sand particles was coated with a thin layer of the wax, i.e. less than 100 nm thick.

### Date Palm Biochar

We produced a nutrient-enriched biochar from the pyrolysis of date palm biomass collected from KAUST Horticultural Department. Dry date palm leaves were crushed using a biomass shredder and tightly packed in batch reactors. The biomass was pyrolyzed in the absence/limited supply of oxygen between 300-500 ℃ for about 3 hrs. After the pyrolysis, we washed the biochar to remove excess salt in the original biomass tissue. The washed biochar was then pre-loaded with macronutrients (Di-ammonium phosphate and micronutrients fertilizers (e.g., Ca, Mg, S, Fe, Mn, Zn, Cu, B, Mo). Through established protocols, we systematically optimized our biochar formulations for enhanced cation and anion exchange capacity, making them able to retain nutrients while slowly releasing the essential macro and micronutrients in fertilizers.

### Plant growth conditions, treatments and experimental designs

We conducted a controlled greenhouse experiment using *M. oleifera* plants at the Plant Growth Core Labs, King Abdullah University of Science and Technology, Saudi Arabia. *Moringa* seeds were sown in plastic trays using potting mix from Stender AG (Schermbeck, Germany). After five weeks, we transplanted the seedlings into pots containing ∼ 4 kg sandy soil watered to two field capacity: 100% field capacity for normal (**N**) irrigation and 50% field capacity for reduced (**R**) irrigation. Weight of each pot was determined gravimetrically. A total of 64 pots were prepared and irrigated 40 pots with plants and 24 pots without plants. Pots without plants were used to quantify water losses due to soil evaporation only. All pots were fertilized by adding 2g/L of NPK (20-20-20) solution with irrigation water on a weekly basis as per the irrigation schedules.

We separated the pots into two groups corresponding to SHS mulch treatment and the controls (herein called bare soil). In each group, half of the pots were subjected to normal irrigation, **N** (100% field capacity) while the other half were maintained under reduced irrigation, **R** (∼50% of normal irrigation). We applied a 10 mm-thick layer of SHS mulch to pots for SHS mulch treatment. Thus, treatment combinations were bare soil-**N**, bare soil-**R,** SHS-**N**, SHS-**R**, biochar-**N**, biochar-**R**, SHS+biochar-**N**, and SHS+biochar-**R**; each with five replicates including pots without plants. All pots were completely randomized giving a 2 × 2 factorial design involving two experimental factors of soil amendment treatment and irrigation regime, each with two levels. Plants were grown in the greenhouse for 25 weeks at 28/19±2 °C (day/night), ∼800 µmolm^-2^ s^-1^ photosynthetically active radiation, and 50-60%±5% relative humidity until harvest.

### Evapotranspiration (ET)

We partitioned ET into soil evaporation and transpiration by performing gravimetric measurements of pot weight on a weekly basis until harvest date. We monitored water loss in pots and adjusted water levels by weighing each pot and adding corresponding amount of water to compensate for the water lost. Weekly ET was taken as total water lost from each pot with plants between the day of irrigation and the next date of weighing the pots using:

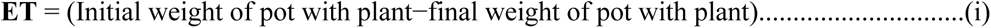

### Evaporation

By assuming that evaporation from pots without plant equals to evaporation from pots with plants, we estimated weekly water loss by evaporation using pots without plants from the expression:

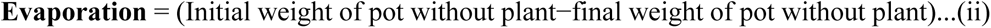

### Transpiration

Weekly transpiration was determined from the expression:

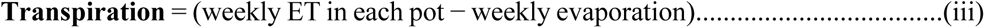

### Plant parameters

#### Plant height

We measured plant height at the beginning of the experiment (i.e., at transplanting) followed by bi-weekly measurements. We monitored changes in plant height over time and calculated the mean plant height at the end of the experiment for each treatment.

#### Leaf Area Index (LAI)

To determine plant leaf area, we took photos from each plant after every two weeks and processed the images. Leaf canopy images were processed using Easy Leaf Area Free software (https://www.quantitative-plant.org/software/easy-leaf-area). We used the RGB value (relative area) of each pixel (i.e., 2 x 2 cm red paper) to identify the leaf and scale regions in each image. Based on this pixel area, we calculated the total area of leaves (canopy size) for each plant. As the RGB scale for green areas remained, the background colors were removed.

#### Trunk diameter

After every two weeks, we measured the trunk diameter of each plant using a digital engineer caliper, to track the increase in girth. We conducted measurements of the trunk around the lower trunk (base height).

#### Stomatal conductance and leaf chlorophyll content

We determined leaf stomatal conductance using AP4 Porometer (Delta T, Cambridge, UK). We performed measurements on three young but fully expanded leaves once a week (between 10:00–12:00) and calculated the mean conductance for each treatment combination. Leaf chlorophyll content index, (CCI) was measured weekly using CCM-200 Chlorophyll Content Meter (Optic-Sciences, Inc. Hudson NH03051, USA), with measurements being performed on four young but fully expanded leaves.

#### Plant biomass

At harvest, we separated the plants into shoot and root components and determined their fresh mass. We then removed leaf samples from each plant for metabolomics profiling, and stored the samples at -80 °C until the time for analysis. We put the other shoot and root samples in paper bags and dried them in an oven at 105 °C for 72 hours after which, their dry mass was determined.

#### Soil sample analysis

At the time of harvest, we collected soil samples from each pot and oven-dried them at 35 ℃ for 7 days before performing soil chemical analysis. To analyze Ca, Mg, Na, K, P, S, Fe, Mn, Cu, and Zn, we weighed 0.5 g of soil sample and extracted using 10 mL of Mehlich 3 solvent25. The mixture was shaken for 4 hours, filtered, and the liquid component was analyzed using Agilent Inductively Coupled Plasma Optical Emission Spectroscopy (ICP-OES). For inorganic nitrogen (ammonium, NH_4_−N and nitrate, NO_3_−N) analysis, 1 g of soil sample was weighed and extracted using 10 mL of 1M KCl, while shaking for 1 hour. The solution was filtered, and the liquid was analyzed using the Salicylate Method for low range nitrogen ammonia (HACH kit no. 2604545) and the 2.6-Dimethylphenol method for nitrogen nitrate (HACH kit no. LCK 339). To analyze pH and TDS, 5 g of the sample was mixed with 10 mL of water for 1 hour. The pH and electrical conductivity (EC) of the resulting slurry (w:v of 1:2) were measured using a SevenCompact Duo with InLab probes.

#### Metabolomics analysis and data processing

For metabolomics analysis, stored leaf samples in vials were immersed in liquid nitrogen and crushed repeatedly using microbeads in the Geno/Grinder^®^ machine at 1200 rpm for 1 min. and the leaves were frozen in liquid nitrogen and lyophilized to dryness. . Cells were later busted using the bead blaster at 4000 rpm for 5 x 1:30 minutes at -10°C. For metabolites extraction, 25mg of each sample was transferred to an Eppendorf vial and 1000µL of 3:1 Methyl-tert-butyl/methanol, followed by shaking at 4 °C for 45 minutes and sonicating in an ice bath for 20 minutes. Subsequently, we added 650µL of 3:1 methanol/water mix with shaking for 1 minute, followed by centrifuging the samples for 5 minutes at 10000 rcf to achieve the phase separation. We then transferred the top organic (MTBE) layer to GC-vials for direct injection in order to test for hydrocarbon contents, while the bottom aqueous layer was transferred to another vial where subjected to complete dryness using Centrivap. After drying, we added 30µL of MOX reagent, followed by rigorous mixing for 1 minute and incubation at 37 °C for 90 minutes at 600 rpm. Subsequently, we added 50µL of the trimethylsilylation (TMS) derevitization reagent i.e., N,Obis(trimethylsilyl)trifluoroacetamide (BSTFA), followed by another incubation at 30 °C for 45 minutes at 600 rpm. Then we performed a non-targeted primary metabolites analysis using a single quadrupole GC-MS (Agilent 7890 GC/5975C MSD) operated under electron ionization (EI) at 70 eV. We analyzed each derivatized sample using both split (20:1) and splitless direct injection into the GCMS inlet. A DB-5MS fused silica capillary column (30 m x 0.25 mm I.D., 0.25-µm film thickness; Agilent J&W Scientific, Folsom, CA) was utilized for chromatographic separation, which was chemically coupled with a 5% phenyl methylpolysiloxane cross-linked stationary phase. Helium was utilized as the carrier gas at a constant flow rate of 1.0 mL min-1. Metabolites were separated using an oven gradient temperature. The initial oven temperature was set to 70 °C for 4 min, then ramped at a rate of 6 °C/min to 330 °C with a 5-min hold time. The GC inlet temperature was set at 250 °C, and the temperature of the transfer line to the MS EI source was kept at 320 °C. The MTBE layer was analyzed through direct split injection, looking for differences in hydrocarbon levels, but we found no significant differences.

We processed the GCMS data using MS-Dial (version 4.9.22121) using a combined MSP library of NIST2020 and GMD (Golm metabolomics Database), freely available online. We performed splitless injection, but due to oversaturation of the sugar region, data regarding peaks with a retention time between 28 and 34 was based on a 1:20 split injection of the same samples. For both, the mass scan range was set at 35-700 Da, with a retention time range of 9-50 min. Minimum peak height was set at 10000. Default deconvolution parameters and identification settings were used. Peak count filter was set at 12.5%, with an N% minimum of 60% per group. Blanks were used to remove background noise and pooled samples were used for alignment in combination with the retention index of an alkane standard. Peak validation was performed in Agilent MSD Productivity ChemStation. Statistical analysis was performed in MetaboAnalyst, comprising the analysis of Variable Importance in Projection (VIP) and Partial Least Square Discriminant Analysis (PLS-DA) scores after normalization by sum and auto-scaling.

### Data analysis

The plant data collected for ET, growth and biomass was analyzed using Origin Pro software (Version 2021). We used a two-way analysis of variance (ANOVA) to analyze the effects of soil amendment and irrigation regime on the variables measured. We subjected all data collected to normality tests; all data conformed to normal distribution requirement. We used Tukey test for 0.05 level of statistical significance to perform multiple comparison of treatment means.

## Results

### Cumulative evapotranspiration (ET), evaporation and transpiration

Our results demonstrate that the highest cumulative ET value attained was 11kg H_2_O lost/plant under N irrigation and the lowest ET attained was 7kg H_2_O lost/plant under R irrigation scenario (**Fig. 1A)**. The cumulative evaporation in SHS mulched-plants under **N** and **R** irrigation was only 28% and 31% of the total ET, respectively (**Fig. 1B**); while transpiration constituted 72% and 69% of the total ET, respectively (**Fig. 1C**). In biochar treatment, cumulative evaporation under either **N** or **R** irrigation was 74% of the cumulative ET while their respective transpiration was 26% of the total ET. For the SHS+biochar treatment, cumulative evaporation was 35.5% and 44.8% of the total ET under **N** and **R**, respectively; thus, cumulative transpiration accounted for 64.5% and 55.2% of the total ET under the respective irrigation regimes. As expected, cumulative evaporation in the control soil was the highest of all, constituting up to 84.7% and 86% of the total ET under **N** and **R**, respectively. Consequently, total cumulative transpiration in control (bare soil) treatment was only 15.3% and 14% of their total ET under **N** and **R** irrigation, respectively.

**Figure 1.**
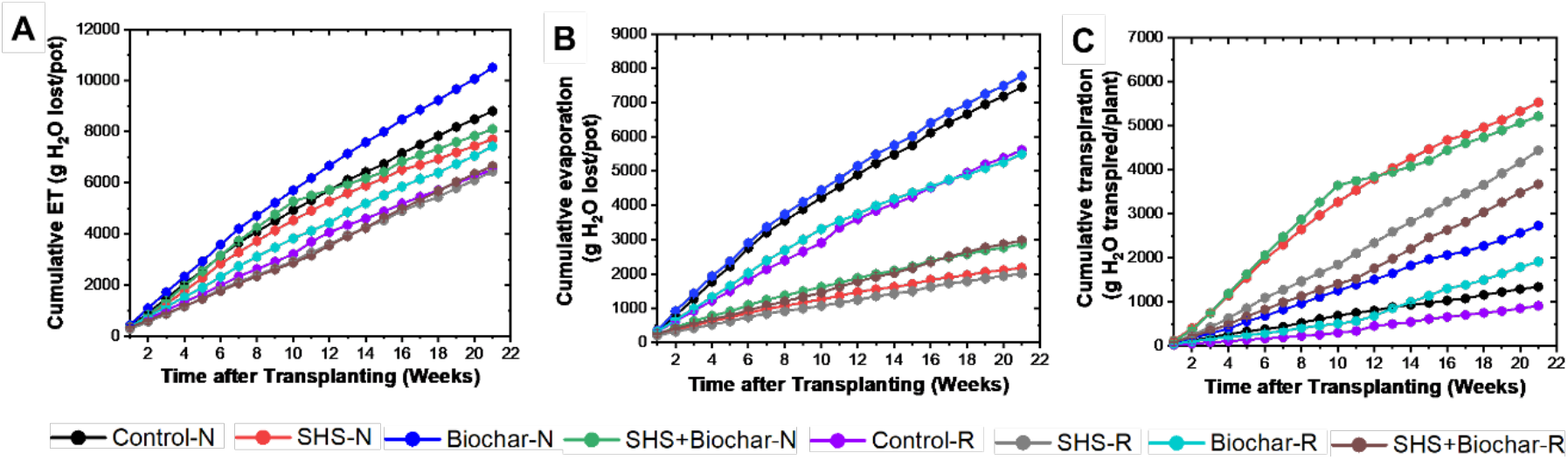
Partitioning of cumulative ET **(A)** into cumulative evaporation **(B)**, and cumulative transpiration **(C)** during 21 weeks for *M. oleifera* plants grown in control soil, Superhydrophobic sand (SHS), biochar, and SHS+biochar treatments under normal **(N)** and reduced irrigation **(R)** during a period of 21 weeks after transplanting. Each data point is a mean of five replicates (*n* = 5).

### Total evaporation and transpiration

Under **N** irrigation, SHS and SHS+biochar significantly reduced total evaporation by 71% and 61%, respectively; both treatments also reduced evaporation by 64% (*p* < 0.05) and 47% (*p* < 0.05), respectively under **R** irrigation (**Fig. 2A**). However, we found no significant difference in evaporation between biochar and control soil under either irrigation regimes. In effect, total transpiration under **N** irrigation significantly increased by 311% and 288% in SHS and SHS+biochar, respectively compared with the controls. Similarly, under **R** irrigation, cumulative transpiration was higher in SHS and SHS+biochar by 385%, and 301%, respectively. Although we found no reduction in evaporative water loss due to biochar treatment, total transpiration was higher than in the controls by 103% and 110% under **N** and **R** irrigation, respectively (*p <* 0.05). When we normalized transpiration values against total leaf area per plant, normalized transpiration values were highest in SHS and SHS+biochar both under **N** (166% and 198%, respectively) and **R** irrigation (142% and 99%, respectively) than in the biochar (50-–54%) and control treatments (**Fig. 2B**) .

**Figure 2.**
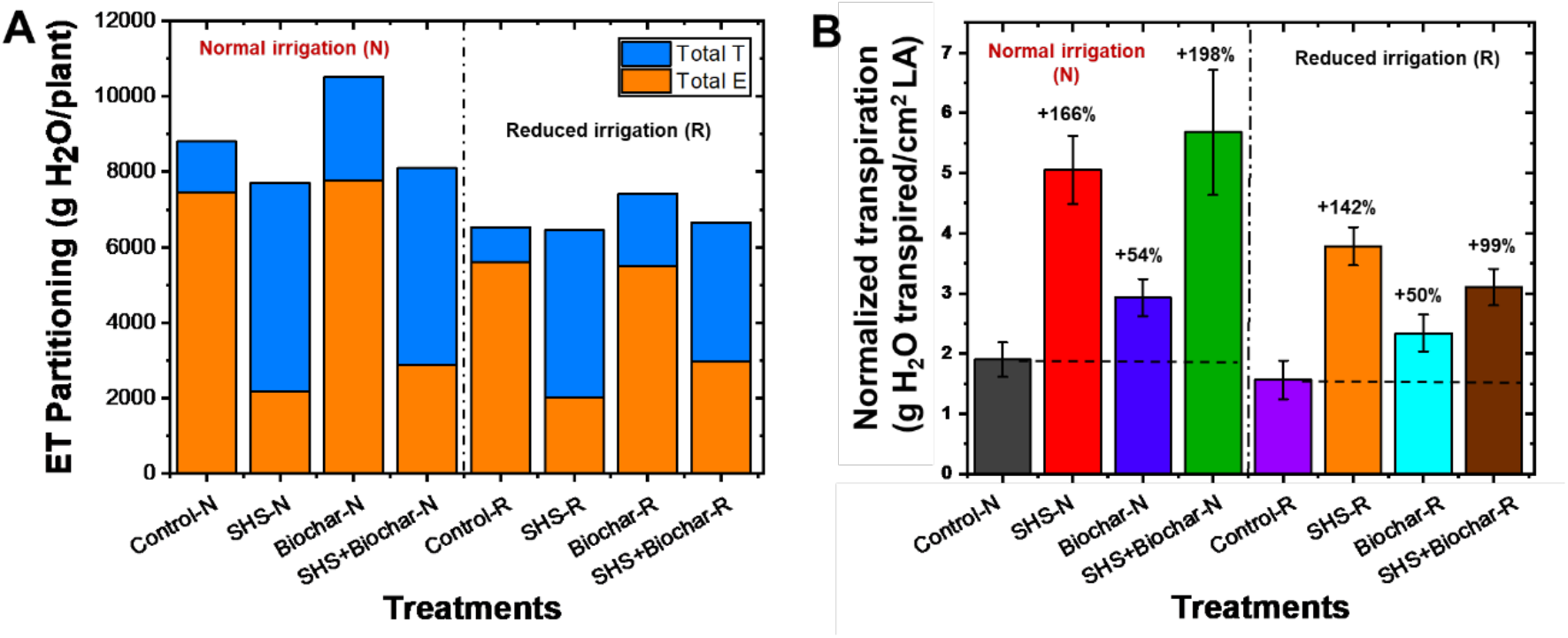
Evapotranspiration (ET) partitioning in Moringa plants. **E** and **T** are total evaporation and transpiration per plant, respectively (**A**) for control (bare) soil, Superhydrophobic sand (SHS), Biochar, and SHS+Biochar treatments under normal **(N)** and reduced irrigation **(R)**. Total transpiration per plant normalized per unit leaf area (LA) in each treatment (**B**); percentage increase in transpiration relative to the control treatment is indicated above each bar. Each data point is a mean of five replicates (*n* = 5).

### Plant growth

Over the growing period, the morphological attributes of the plants from different treatments were visually compared side-by-side. **Fig. 3A** shows the representative snapshots of the plants at selected weeks during their growth period. Generally, plants grown with SHS, Biochar, and SHS+Biochar grew taller and bigger in size than their control counterparts. Meanwhile, plants grown with biochar alone grew less than those grown with SHS or SHS+Biochar.

**Figure 3.**
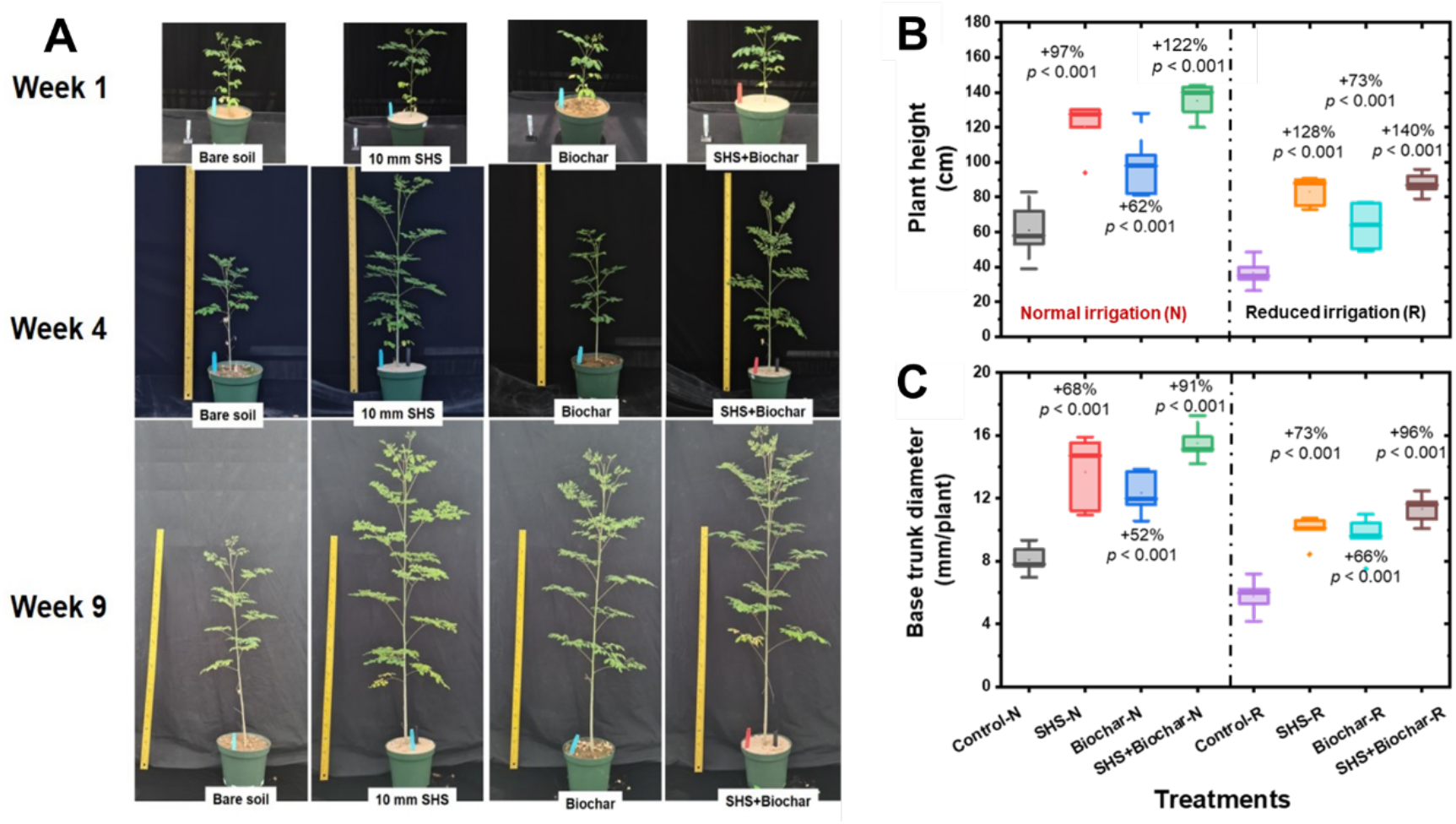
**A.** *Moringa* plants growing in the greenhouse using bare (control) soil, Superhydrophobic sand (SHS) mulch, Biochar, and SHS+Biochar combination under normal (**N**) irrigation at 9 weeks after transplanting. Boxplots showing **B-** mean final plant height; and **C-** mean base trunk diameter for plants grown in control (bare) soil, SHS, biochar, and SHS+Biochar under normal (**N**) and reduced (**R**) irrigation scenarios at 21 weeks after transplanting. Each box represents the data distribution from 5 replicates (*n* = 5; *N*=40) with the mid-line indicating the median value, the white dot inside the box represents the mean value, the upper and lower sections of the box represent the 25% and 75% of data points, respectively, and the whiskers on the box represent the 1.5 interquartile range, the dot outside the box indicates outlier. Percentage differences between treatments are presented relative to control (bare) soil along with their corresponding *p*- values derived from two-way ANOVA using *p* < 0.05 level of statistical significance.

### Plant height and trunk diameter

Under **N** irrigation, mean plant height significantly increased in SHS, biochar and SHS+biochar plants by 97%, 62%, and 122%, respectively relative to the controls (**Fig.3B**). Under **R** irrigation, mean plant height was higher in SHS, biochar, and SHS+biochar than their control counterparts by 128%, 73%, and 140%, respectively. The base trunk diameter also increased significantly due to SHS, biochar, and SHS+biochar treatments by 68%, 52%, and 91% when **N** irrigation was used (**Fig. 3C**). In the same trend, base trunk diameter increased under **R** irrigation by 73%, 66%, and 96% for SHS, biochar, and SHS+biochar treatments in comparison with the control plants (*p* < 0.05).

### Leaf Area Index

From the visual depiction in **Fig. 4A**, canopy sizes for plants grown in SHS and SHS+biochar were significantly larger than in their bare (control) soil counterparts, except for biochar treatment, which did not show significant difference from the control plants under either **N** or **R** irrigation. In fact, compared with bare soil plants, leaf area in SHS and SHS+biochar was significantly larger under **N** irrigation by 57% and 132%, respectively; followed by 145% and 124% under **R** irrigation, respectively (**Fig. 4B**).

**Figure 4.**
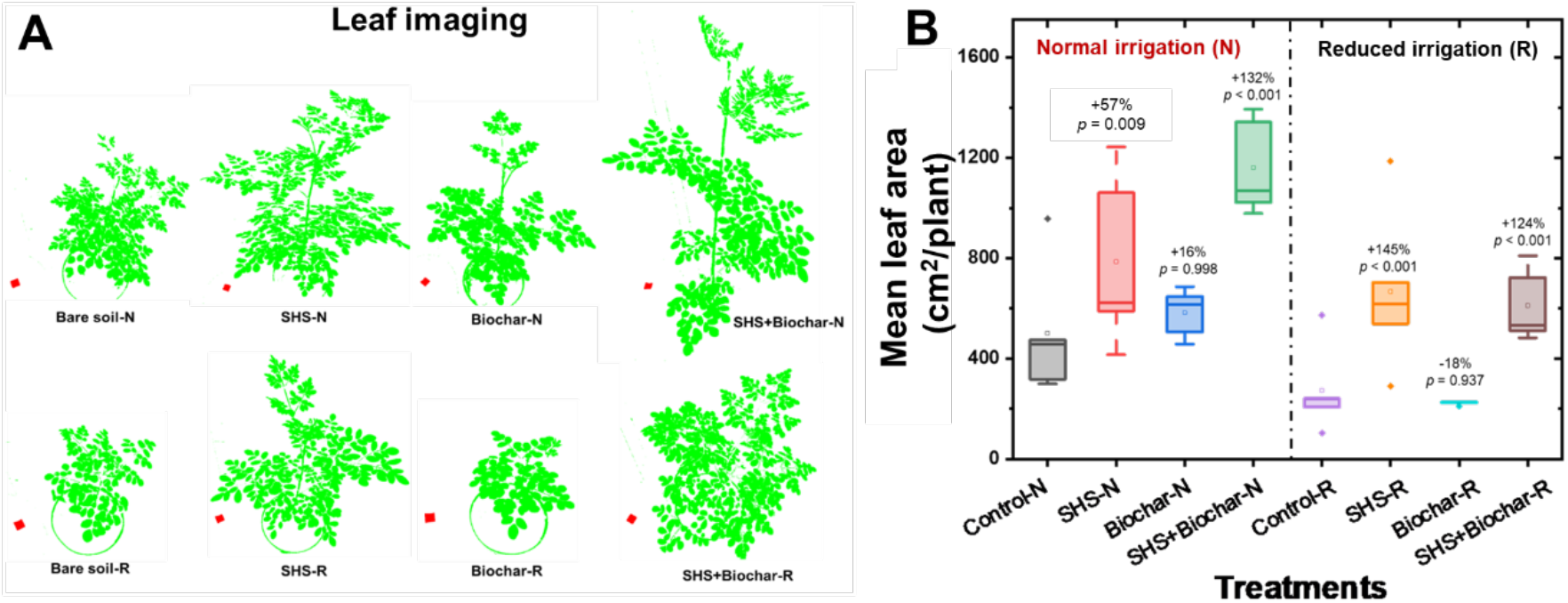
Measurement of leaf area. Processed images of plant leaf canopies (**A**) and mean leaf area (**B**) of *Moringa* plants grown in Bare soil, SHS, Biochar, and SHS+Biochar under normal (**N**) and reduced (**R**) irrigation. Leaf canopy images were processed using Easy Leaf Area Free software (https://www.quantitative-plant.org/software/easy-leaf-area). We used the RGB value (relative area) of each pixel (i.e., 2 x 2 cm red paper) to identify the leaf and scale regions in each image. As the RGB scale for green areas remained, the background colors were discriminated.

### Leaf chlorophyll content (CCI) and stomatal conductance

Our results show that, relative to bare soils, total CCI significantly increased in SHS, biochar, and SHS+biochar by 11%, 14%, and 13%, respectively under **N** irrigation; and by 19% and 11% under **R** irrigation for SHS and SHS+biochar treatments, respectively (**Fig.5A**). However, we did not find significant increase in CCI in biochar under **N** or **R** irrigation (*p* > 0.05). Under N irrigation, leaf stomatal conductance increased in SHS, biochar and SHS+biochar by 131%, 51%, and 175% (*p* < 0.001), respectively relative to the bare soils (**Fig. 5B**). Under **R** irrigation, however, biochar treatment showed significantly lower stomatal conductance (16%) than the control, whereas conductance in SHS and SHS+biochar significantly increased by 63% and 82%, respectively.

**Figure 5.**
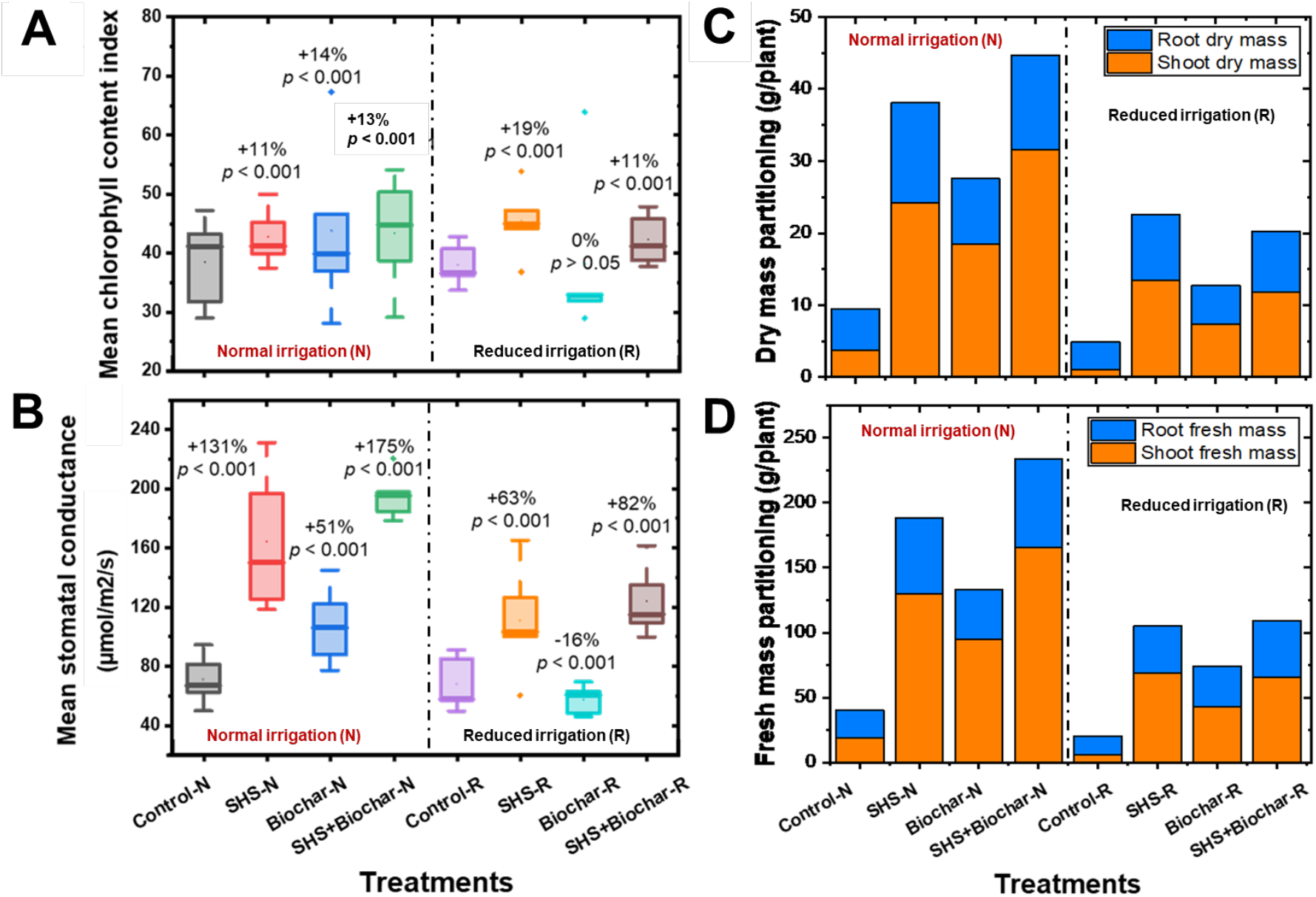
Boxplots showing **A-** mean chlorophyll content index; **B-** mean stomatal conductance, for plants grown in control (bare) soil, SHS, biochar, and SHS+Biochar under normal (**N**) and reduced (**R**) irrigation scenarios. Biomass distribution in *Moringa* plants under different soil treatments and irrigation regimes: **C-** shoot and root dry mass partitioning, **D-** shoot and root fresh mass partitioning in different treatments under normal and reduced irrigation. Each box/bar represents the data from 5 samples (*n* = 5). For the box plot, the mid-line indicates the median value, the white dot inside the box represents the mean value, the upper and lower sections of the box represent the 25% and 75% of data points, respectively, and the whiskers on the box represent the 1.5 interquartile range, the dot outside the box indicates outlier. Percentage differences between treatments are presented relative to control (bare) soil along with their corresponding *p*-values derived from two-way ANOVA using *p* < 0.05 level of statistical significance.

### Biomass partitioning

All soil amendment treatments had profound effects on both dry and fresh shoot biomass increases than on root biomass (**Fig. 5C & D**). Under **N** irrigation, shoot dry mass increased due to SHS, biochar and SHS+biochar treatments by 542%, 390%, and 738%, respectively in comparison with the control. Meanwhile, shoot dry mass increased under **R** irrigation by 1271%, 648%, and 1102% in SHS, biochar and SHS+biochar treatments, respectively (*p <* 0.005). For roots, dry mass increased under **N** irrigation due to SHS, biochar and SHS+biochar by 142%, 52% and 129%, respectively (*p <* 0.05); also, root dry mass increased under **R** irrigation in SHS, biochar and SHS+biochar treatments by 133%, 37%, and 115%, respectively (*p <* 0.05).

### Metabolites analyses

Analyses of *Moringa* leaf tissue showed significant variation in metabolite profiles for plants in the control versus those in SHS, biochar, or their combination, under **N** and **R** irrigation (**Fig. 6**). In **Fig. 6A**, we present the top 40 metabolites contributing to the variation of metabolite profiles under the different treatment conditions based on the Variable Importance in Projection (VIP) scores. In this case, we considered that metabolites with a VIP score of ≥ 1.0 are of significant importance in contributing to the differences between treatment groups (i.e., **19** metabolites involved ranging from Homoserine to L-Aspartic acid). From the colored scale (red to blue), our results demonstrate that leaves of plants in the control treatment had the highest concentration or abundance of 13 amino acids (i.e., from Homoserine to L-Tryptophan, L-Methionine, L-5- Oxoproline, L-Aspartic acid) and the nucleoside, guanosine. Such metabolite abundance in control plants relative to other treatments apparently followed a similar pattern under both **N** and **R** irrigation, except for L-5-Oxoproline and L-Aspartic acid. For VIP scores > 1.0, SHS and biochar had the highest relative abundance of 3 sugars (i.e., D-Mannose, D-Fructose, glucose) and one organic acid (i.e., malic acid) under both irrigation regimes.At VIP scores < 1.0, the relative abundance of sugars (such as sucrose to L-(-)-Arabitol)), fatty acid (palmitic acid), organic acid (succinic acid), and nucleoside (adenosine) was generally higher for plants in control and biochar treatments under both **N** and **R** irrigation.

**Figure 6.**
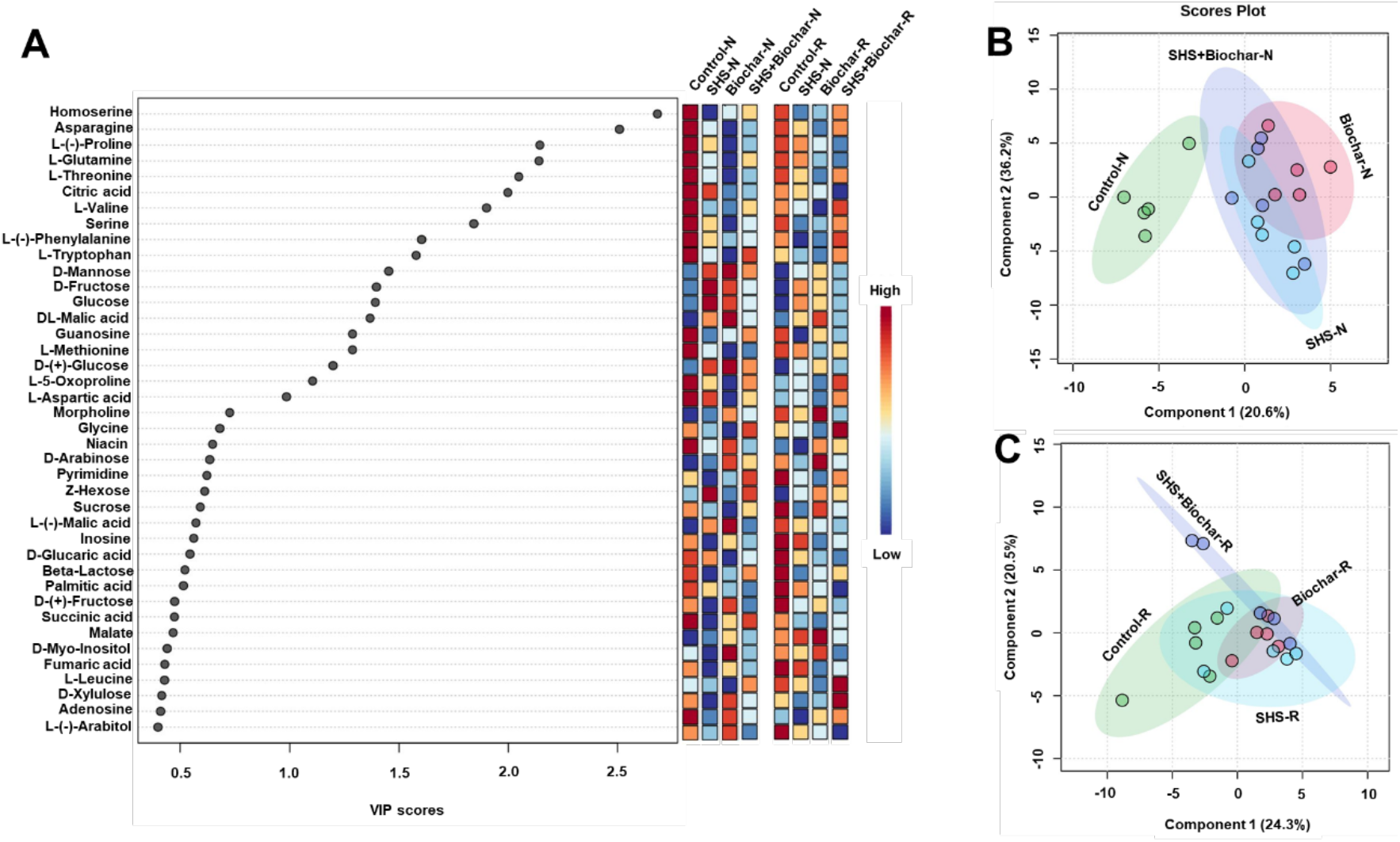
Variable Importance in Projection (VIP) scores for the top 40 metabolites contributing to variation in metabolic profiles of controls vs. SandX (SHS), Biochar, and SHS+Biochar treatments under normal (**N**) and reduced (**R**) irrigation **(A)**. The colored scale indicates the relative abundance of metabolites ranging from **RED** to **BLUE** representing the low and high, respectively. Partial least Square-discriminant analysis (PLS-DA) scores plots of significant metabolites in ANOVA derived for SHS, Biochar, SHS+Biochar vs the controls under **N** irrigation **(B)** and R irrigation **(C)**.

Further analysis using partial least square discriminant analysis (PLS-DA) to provide insights into the metabolic profiles of the plants under different treatments and irrigation rendered a two-component model (**Fig. 6B-C**) as calculated by cross-validation (CV). The PLS-DA scores discriminated the control treatment from SHS, Biochar, and SHS+Biochar samples into four obvious clusters separated by scores of the first component; the metabolite profile of control samples mostly positioned towards the negative end of component 1 while those of SHS, Biochar and SHS+Biochar are mainly positioned at the positive side of component 1 under both irrigation regimes.

### Soil chemical properties

Analysis of soil properties revealed that pH of the control soil did not significantly differ before and at the end of the experiment (**Table 1**). Final analysis demonstrates significant increase in the soil pH due to biochar from 7.6 in no-biochar treatments (i.e., control and SHS) to about 8.6 in the biochar-amended soils (i.e., Biochar, SHS+Biochar). Compared with the control soil, both the EC and salinity for SHS, biochar and SHS+Biochar-treated soils significantly reduced by 3, 4, and 17 times, respectively under **N** irrigation; soil EC and salinity reduced by 3, 3, and 12 times, respectively under **R** irrigation. On the other hand, under **N** irrigation, total nitrogen (NH_4_–N+NO_3_–N) in SHS, Biochar, and SHS+Biochar-amended soil was less than in the control soil by 2, 3, and 12 times less, respectively. Under **R** irrigation, total nitrogen concentration in SHS, Biochar, and SHS+Biochar was respectively 3, 3, and 6 times less than in the control. Phosphorus concentration in both SHS and biochar was 2-times more than in the control under **N** irrigation while SHS+Biochar did not significantly differ from the control. Under **R** irrigation, phosphorus concentrations from SHS, Biochar and SHS+Biochar-treated soils were 2, 6, and 6-times more than in the control (p < 0.05). While the concentration of potassium did not change between SHS and control soils, biochar and SHS+Biochar-treated soils had 2 and 4 times less concentration of potassium under **N** irrigation. Under R irrigation, the concentrations of potassium in SHS and biochar were higher than in the control soils by 19% and 26%, respectively; while SHS+Biochar-soil did not significantly differ from the control soil.

**Table 1.** Soil chemical properties before and after the experiment. Changes in soil pH, electrical conductivity (EC), salinity, elemental compositions and inorganic nitrogen (NH_4_–N and NO_3_–N) concentrations in control soil, SandX (SHS), Biochar, SHS+Biochar under normal (**N**) and reduced (**R**) irrigation.

| Treatments | Soil pH, EC, Salinity |  |  | Elemental Concentrations (ppm) |  |  |  |  |  |  |  |  |  |  | Inorganic N Content |  |
| --- | --- | --- | --- | --- | --- | --- | --- | --- | --- | --- | --- | --- | --- | --- | --- | --- |
|  | pH<br>(1:2, w/v) | EC<br>(μS/cm) | Salt<br>(ppm) | Al | Ca | Cu | Fe | K | Mg | Mn | Na | P | S | Zn | NH<sub>4</sub>-N<br>(mg/kg) | NO<sub>3</sub>-N<br>(mg/kg) |
| Original soil | 7.9 | 420 | - | 624 | 3237 | 1.25 | 286 | 220 | 617 | 39.1 | 414 | 42.2 | 51.2 | 2.32 | 0.37±0.02 | 16.2±0.91 |
| Control-N | 7.7 | 2631 | 3368 | 578 | 3410 | 9.77 | 248 | 812 | 679 | 45.6 | 758 | 142 | 80.8 | 12.9 | 0.52±0.19 | 7.61±0.70 |
| SHS-N | 7.5 | 769 | 986 | 535 | 2870 | 2.81 | 214 | 726 | 560 | 43.3 | 596 | 232 | 44.4 | 17.2 | 0.54±0.13 | 4.41±0.82 |
| Biochar-N | 8.4 | 679 | 869 | 486 | 3104 | 3.52 | 188 | 357 | 567 | 44.7 | 733 | 230 | 54.6 | 15.8 | 0.23±0.04 | 2.42±0.59 |
| SHS+Biochar-N | 8.8 | 157 | 201 | 456 | 2381 | 2.50 | 196 | 206 | 483 | 36.0 | 518 | 187 | 42.3 | 12.4 | 0.25±0.02 | 0.41±0.06 |
| Control-R | 7.8 | 1792 | 2294 | 283 | 1449 | 0.95 | 126 | 280 | 297 | 20.1 | 312 | 39.1 | 28.6 | 6.23 | 0.92±0.35 | 7.17±0.31 |
| SHS-R | 8.4 | 697 | 892 | 587 | 3080 | 1.57 | 247 | 346 | 593 | 42.2 | 622 | 63.3 | 56.9 | 12.9 | 0.55±0.38 | 2.45±0.40 |
| Biochar-R | 8.4 | 596 | 763 | 581 | 2991 | 3.93 | 246 | 383 | 647 | 50.1 | 699 | 224 | 62.2 | 14.5 | 0.20±0.05 | 2.36±0.11 |
| SHS+Biochar-R | 8.6 | 148 | 189 | 501 | 2830 | 3.27 | 217 | 252 | 537 | 40.4 | 535 | 215 | 50.3 | 12.4 | 0.26±0.02 | 1.02±0.67 |

## DISCUSSION

Here, we discuss our results in terms of the mechanisms behind the observed responses of *M. oleifera* to SHS mulch and biochar application under the two contrasting irrigation scenarios. We explain the observed results on the plant’s ET partitioning and their relationships with the plant growth (morpho-physiological) parameters, as well as the results on metabolic profiling and changes in soil chemical properties.

On account of its extreme water repellency^23, 25^, our SHS significantly reduced evaporation of water from the soil and substantially contributed to enhanced soil water retention. The water retained in the soil due to the suppressed evaporation gets channeled towards transpiration by the plant’s vascular systems^24, 26^, as was demonstrated by the significant increases in transpiration due to the SHS and SHS+biochar treatments under **N** or **R** irrigation (**Fig. 1-2**). This amount of water transpired in *Moringa* due to SHS application is of great importance as it is associated with growth and yield improvement following the process of photosynthesis^26–28^. Although evaporation in biochar treatment was as high as in the control, total transpiration in biochar alone was about two-times higher than in the control treatment. The nutrient-enriched biochar could have enhanced nutrient-availability and its use efficiency for biomass development, which even raised the plant’s transpiration demand higher than in the control plants.

We attribute the high evaporation observed in our biochar-amended soil to biochar properties that affect its water holding capacity and make the soil susceptible to evaporation. For instance, biochar hydrophobicity (i.e. its water-repelling property) influences biochar’s water holding capacity due to the development of hydrophobic functional groups in the biochar surface during biomass pyrolysis^29, 30^. In a study, all biochar types tested were mostly hydrophobic due to less polar components on their surface, in which case such biochar could exhibit reduced water retention efficiencies^31^. This water-repellent hindrance to water retention of such a biochar could explain why our SHS+Biochar treatment did not show additive synergistic benefits relative to the individual effects of SHS and biochar (**Fig. 1B & C; 2A**).

Our study reveals that by reducing soil evaporation using SHS mulch, we optimized the amount of water retained in the soil and increased the efficiency of transpiration towards facilitating plant growth and biomass production^26^. In the same light, the enriched biochar provided a synergy for improved nutrient usage, thereby, promoting growth, photosynthesis and biomass formation^32, 33^. These effects were demonstrated via the increased plant height and base trunk diameter (**Fig. 3**), leaf area (**Fig. 4**), leaf chlorophyll content index, stomatal conductance, as well as the fresh and dry biomass of shoots and roots (**Fig. 5**) observed in the SHS and biochar treatments, as well as in their combination. In related studies, plastic mulching significantly improved flowering and panicle number in *M. oleifera* plants^34^; while *Gliricidia* biochar also showed significant positive effects on the growth of Moringa in terms of plant height, stem diameter, dry matter yield and root: shoot ratio^35^. Consistent with our findings under SHS mulching, soil water availability positively correlates with leaf area, stomatal conductance, transpiration, and biomass^24, 36^.

Following metabolomics profiling, the high relative abundance of most amino acids, sugars, fatty acid, and organic acid shown in the leaves of control plants relative to those in other treatments could be attributed to the plants’ physiological response to water or nutrient stress^37^ induced by excessive water loss via evaporation. The increased accumulation of amino acids such as proline, L-Tryptophan, L-Phenylalanine, Aspartate, etc. (**shown in Fig. 6A**) occurs in response to drought stress^38–40^. These amino acids play an important role in maintaining cellular integrity during stress and help in plant recovery from and adaptability to environmental stresses^41–43^. For instance, proline acts as an osmolyte for osmotic adjustment, and contributes to stabilizing sub-cellular structures (e.g., membranes and proteins), scavenging free radicals and buffering cellular redox potential under stress conditions^44^. Alongside its role in normal development, asparagine has been shown to accumulate in response to water stress, and nutrient deficiencies^45^. Meanwhile, the high concentration of D-Mannose, D-Fructose, glucose, and malic acid in plants grown with SHS or biochar treatment under **N** and **R** irrigation can be attributed to the increased water/nutrient-use efficiency and carbon assimilation owing to higher rates of photosynthesis than plants in control soils.

The positive effect of biochar has been attributed to the combination of the biochar and nutrients in fertilizer and associated soil microbes, rather than the fertilizer effect alone^33^. This observation underscores the significant role of biochar in soil nutrient- buffering and providing conducive microsites/pores for beneficial microbes^17^. Although microbial analysis was beyond the scope of this work, our results show that the adsorptive property of biochar improves soil physical and chemical properties^46, 47^, as shown by the increase in soil pH and composition of essential elements (**Table 1**). Our results suggest a higher nitrogen use-efficiency by plants in biochar-amended soils (Biochar and SHS+Biochar) as demonstrated by the lower concentrations of NH_4_-N and NO_3_-N, corroborated by the higher biomass produced in biochar-amended soils than in the control soils. Similarly, the low concentrations of potassium in biochar-amended soils under **N** irrigation suggests that a higher amount of potassium in the soils was taken up and assimilated by the plants than those in the control case. From our results, the high final concentration of phosphorus found in biochar-amended soils support the high P adsorption capacity of biochar, hence, the application of phosphorus-rich biochar as a soil amendment or a slow-release phosphorus fertilizer to promote plant growth^48^.

Hence, by pre-loading our date palm biochar with nutrients, we capitalized on the porous and cationic exchange property of the biochar for enhanced nutrient retention and release as required by the plants. The positive effect of SHS and enriched biochar on the growth of *Moringa* suggests that date palm biochar could play an important role in addressing the challenges involved in arid land rehabilitation through greening or afforestation initiatives.

## CONCLUSION

We investigated the effects of SHS mulching and enriched date palm biochar, and their combination on the ET and growth of *M. oleifera* under normal and reduced irrigation scenarios in a greenhouse. By reducing soil evaporation, SHS mulch maximized the amount of water retained in the soil and increased the efficiency of transpiration towards facilitating plant growth and biomass production. In the same light, the enriched biochar provided a synergy for improved nutrient usage, thereby, promoting growth, photosynthesis and biomass formation. Moringa plants grown in unamended (control) soils accumulated higher concentrations of osmolytes such as amino acids, sugars, nucleosides, organic acids and fatty acids than those of other treatments in response to water or nutrient stress. High water/nutrient use efficiency and carbon assimilation in plants under SHS and biochar is associated with increased accumulation of certain sugars and organic acids such as D-Mannose, D-Fructose, glucose, and malic acid. The positive effects demonstrated by SHS and date palm biochar on Moringa plant underscore the potential of these complementary technologies in addressing the challenges of water scarcity and nutrient deficiency in arid land for promoting both crop production and desert revegetation in Saudi Arabia and the Middle East at large.

## Acknowledgments

We acknowledge the support of the following individuals towards the success of this study:

● Prof. Magdi A. Mousa – assistance during our initial testing of our batch-scale biochar reactor at King Abdulaziz University field site (Hada Al Sham),

● The KAUST Plant Growth Core Labs Team – Greenhouse facilities support

● Rafa M. Alyubi and Wafa Taeif– visiting students/interns for their assistance during the experiment.

## Author Contributions

**KO:** Conceptualization, experimental design and execution; ET and plant data collection and analysis, manuscript writing

**BV:** Sample preparation for metabolomics profiling, GC-MS analysis, and metabolomics data processing.

**BA:** Biochar treatment and ET data collection

**LOE:** Irrigation and ET data collection, leaf area index and imaging

**AA:** Irrigation and plant monitoring, ET data collection and imaging

**NVHM:** Performed soil analyses (pH, EC, salinity, ICP-OES elemental composition, NH4-N/NO3-N).

**AAGH**: Reactor design and testing, date palm biochar production and post-processing.

**AGJ:** Biochar production and treatment, manuscript review/revision

**NK:** Performed metabolomics profiling and oversaw on all analytical steps: from samples preparation, GC-MS injection and data processing.

**HM:** Conceptualization, supervision, review and editing, supervision, funding and resource acquisition.

## Funding

This work was supported with funds from King Abdullah University of Science and Technology (KAUST) through the Baseline Funding to HM (# BAS/1/1070-0101)

## Data availability statement

The authors declare that all the data supporting the findings of this study are available within the paper.

## Conflicts of Interest

HM and AGJ were awarded a patent **US20200253138A1** for the SHS material.

## Notes

### Competing Interest Statement

Himanshu Mishra and Adair Gallo Jr. were awarded a patent US20200253138A1 for the engineered Superhydrophobic Sand Mulch Technology

